# Selective chr21 homolog silencing reveals polymorphisms influence the epigenetic silencing and functional dosage of RWDD2B

**DOI:** 10.1101/2025.09.17.676855

**Authors:** Eric C. Larsen, Jennifer E. Moon, Oliver D. King, Jeanne B. Lawrence

**Affiliations:** Department of Neurology, University of Massachusetts Chan Medical School, Worcester, MA, United States 01605; Department of Pediatrics, University of Massachusetts Chan Medical School, Worcester, MA, United States 01605

## Abstract

Polymorphisms that affect chr21 gene expression have significance for both variable severity in Down syndrome and common multifactorial conditions. Results here demonstrate “selective homolog silencing” in cells from even one individual can provide a valuable complement to large studies. In trisomic iPSC subclones that silence different chr21 homologs (via *XIST*-based silencing), we discovered unusually large, homolog-specific, differences in *RWDD2B* in iPSCs, cortical organoids and endothelial cells. RNA FISH showed *RWDD2B* transcription almost entirely from the H1 homolog, correlated with CpG promoter methylation differences. Polymorphisms different on H1 versus H2/H3 had strongest eQTLs in GTEx, especially in brain. Collective results indicate RWDD2B functional dosage is more frequently disconnected from copy number even compared to neighboring genes. RWDD2B function is unknown, but nearby methyl-eQTLs are implicated in osteoarthritis, and potential roles in inflammation or immune response merit consideration. This study has significance for RWDD2B regulation and demonstrates a cell-based methodology to study polymorphisms.

## Introduction

Down syndrome (DS/T21), caused by trisomy of chromosome 21 (chr21), remains a leading cause of intellectual disability, occurring in ∼1/700 live births in the U.S. While children with DS are typically sociable and happy, they often face various medical challenges. Consistent characteristics vary in severity but include distinct craniofacial features, short stature, cognitive disability, vulnerability to respiratory infection, and accelerated aging, with an 80% occurrence of early-onset Alzheimer’s disease (AD). In addition, T21 greatly elevates risks for congenital heart defects, transient myeloproliferative disorder (TMD) and leukemia, and moderately elevates risk for many common multifactorial conditions including autoimmune conditions, autism, arthritis, muscle hypotonia, and orthopedic issues such as hip dysplasia^1–5^. While triplication of the *APP* (amyloid precursor protein) gene on chr21 in DS was critical to identifying APP’s role in AD, some evidence suggests that other chr21 genes may reduce penetrance of *APP* triplication^6^. Hence, understanding how T21 influences risks for various conditions has relevance to the broader population. The cellular mechanisms that lead to multi-systemic effects in DS are not well understood, nor is it known how many of the ∼250 coding chr21 genes are dosage-sensitive and contribute to pathology when present in three copies. This is more challenging for conditions like chronic inflammation and autoimmunity, which are of growing importance in DS, as multiple chr21 genes are known to be involved^7–10^. Consistent characteristics will be due to having three copies of chr21 genes, but variable severity and co-morbidities will depend on genetic polymorphisms on or off chr21^11–13^. Hence, identifying polymorphisms that modify expression levels (i.e. functional dosage) has enhanced importance for DS since dosage-sensitive chr21 genes are already at or near pathological thresholds.

Genome-wide association studies (GWAS) have identified specific DNA loci—known as quantitative trait loci (QTLs)—where polymorphisms are associated with particular traits or phenotypes^14,15^. Among these, expression QTLs (eQTLs), which influence gene expression levels, are most extensively studied^16^. QTLs can also affect other molecular processes such as DNA methylation (mQTLs)^17^, RNA splicing^18,19^, RNA stability^20,21^, and translation rates^22,23^, which may or may not alter gene expression levels. When a QTL associated with DNA methylation is moreover associated with gene expression levels, it is referred to as a methylation-expression QTL (meQTL)^24–27^. These typically only modestly modulate expression levels, unlike the binary “on-off” expression (i.e. transcriptional silencing) associated with promoter methylation in developmental processes such as X-chromosome inactivation in early embryogenesis or gene imprinting during gametogenesis^28,29^.

The Genotype-Tissue Expression (GTEx) program (and similar studies) examines samples from many individuals with corresponding genomic and transcriptomic data to identify eQTLs genome-wide. To minimize the confounding effects of broad genomic variation between individuals, eQTLs from GTEx are based on tissue samples from ∼1,000 individuals and leverages natural recombination to narrow eQTL locations. Similarly, other recent efforts to use cell-based approaches to study eQTLs have also required samples from hundreds of individuals^30–33^. Adding to the challenge, eQTLs typically have weak or modest effects on expression, and some effects only occur if several variants co-occur^34,35^. As GTEx examines eQTLs in different tissues, which contain multiple cell-types, more methodologies are needed that can examine specific cell types, since eQTLs can be cell-type specific^16^. In addition, better ways are needed to understand how polymorphisms impact cell development and function to bridge the gap between QTLs and phenotypic traits identified in GWAS^36,37^. Differentiation of induced pluripotent stem cell (iPSC) lines is a possible approach, but the culture and differentiation of hundreds of iPSC lines is highly laborious and costly, making this less practical^30,32,33^.

Here we illustrate an approach that can help circumvent some of the challenge, by manipulating the haplotype expressed from part of the genome while keeping the rest of genomic background constant in cells from a single individual. This is shown in T21 cells using selective silencing of one chr21 copy, which represents just 1% of the genome. This is an extension of our lab’s system for *XIST*-based trisomy silencing to focus on “selective homolog silencing”. Our lab previously demonstrated the concept of trisomy silencing by inserting an inducible *XIST* transgene into one chr21, in iPSCs derived from an individual with DS, where *XIST* transgene induction comprehensively silences the extra chr21^38^. This allowed for direct comparison of cultures of the same cell population with or without increased chr21 dosage, which we have used to demonstrate impacts of T21 in various iPSC-derived cell-types, identifying potential pathways of pathology^38–41^.

Initially the *XIST* transgene inserted randomly into different chr21 homologs^38^, which we capitalize on here to compare the effects of silencing different chr21 homologs. This strategy led to the discovery of an unusually strong effect of *cis* polymorphisms on *RWDD2B* expression. By examining impacts of selective homolog silencing, we demonstrate a particular homolog primarily transcribes *RWDD2B* (H1^RWD^), and that CpG promoter methylation correlates with *RWDD2B* transcriptional status. By cross-referencing with eQTLs from GTEx we found variants different on H1^RWD^ correspond with increased *RWDD2B* expression, with large effect size, while those different on H2/H3 correspond with lower expression and smaller effect size, consistent with genetic variants underly this difference. Furthermore, public data on cells derived from other individuals (with or without T21) also show large variation in *RWDD2B* expression levels between individuals. Importantly, our data from cortical organoids is consistent with GTEx data that shows eQTL effects are strongest in brain regions. *RWDD2B* has received far less attention in DS than many chr21 genes, but an earlier zebrafish study showed *RWDD2B* over-expression produced a phenotype^42,43^. While the gene’s function is unknown, a recent study identified RWDD2B as the top target of TRIM7 for proteomic degredation^44^, of potential interest in DS, and *RWDD2B* meQTLs have been implicated for osteoarthritis (OA) risk^26,45,46^ (see Discussion). Interestingly, there is growing evidence to suggest both the importance of chronic inflammation^47–49^ and epigenetics^50,51^ in OA. Thus, chronic inflammation is of growing interest in both DS and OA.

While we focus here on *RWDD2B*, the concept of selective haplotype silencing, while keeping genomic background constant, has broader potential as a complement to other methodologies. We briefly discuss and illustrate how expansion of this approach can enable identification of other candidate chr21 eQTLs, and, importantly, facilitate the study of cell-type specific and downstream effects of polymorphisms.

## Results

This study utilizes a panel of 10 isogenic subclones of induced pluripotent stem cells (iPSCs) isolated from a single iPSC line derived from an individual with Down Syndrome (DS/T21), detailed in Figure 1A. When we engineered the original parental line (par) with a TET3G inducible *XIST* transgene, we isolated several *XIST*-transgenic subclones (c1, c2, c3, c5, c5a, and c7 utilized in this study) as well as both trisomic (parA, parB) and disomic (dis, disA, disB) subclones without the *XIST* transgene^38^. In several recent studies using this cell panel to investigate effects of trisomy for chr21, we examined three to four *XIST*-transgenic clones with the expectation that core (consistent) changes due to T21 should be similar across clones. These prior studies showed *XIST* RNA comprehensively silences genes across chr21. While it had previously been suggested there would be substantial feedback regulation to reduce chr21 over-expression, studies from our lab and others^40,41,52^ find that chr21 mRNA levels generally approximate the expected 1/3 reduction with silencing or loss of one chr21 (Fig 1B + Fig S1A+B, Tables S1-S3). While modest deviations from theoretical expectations may be meaningful and are discussed later, one chr21 gene showed unusually large fold-changes in our data, which led us to investigate the basis for this further.

**Figure 1:**
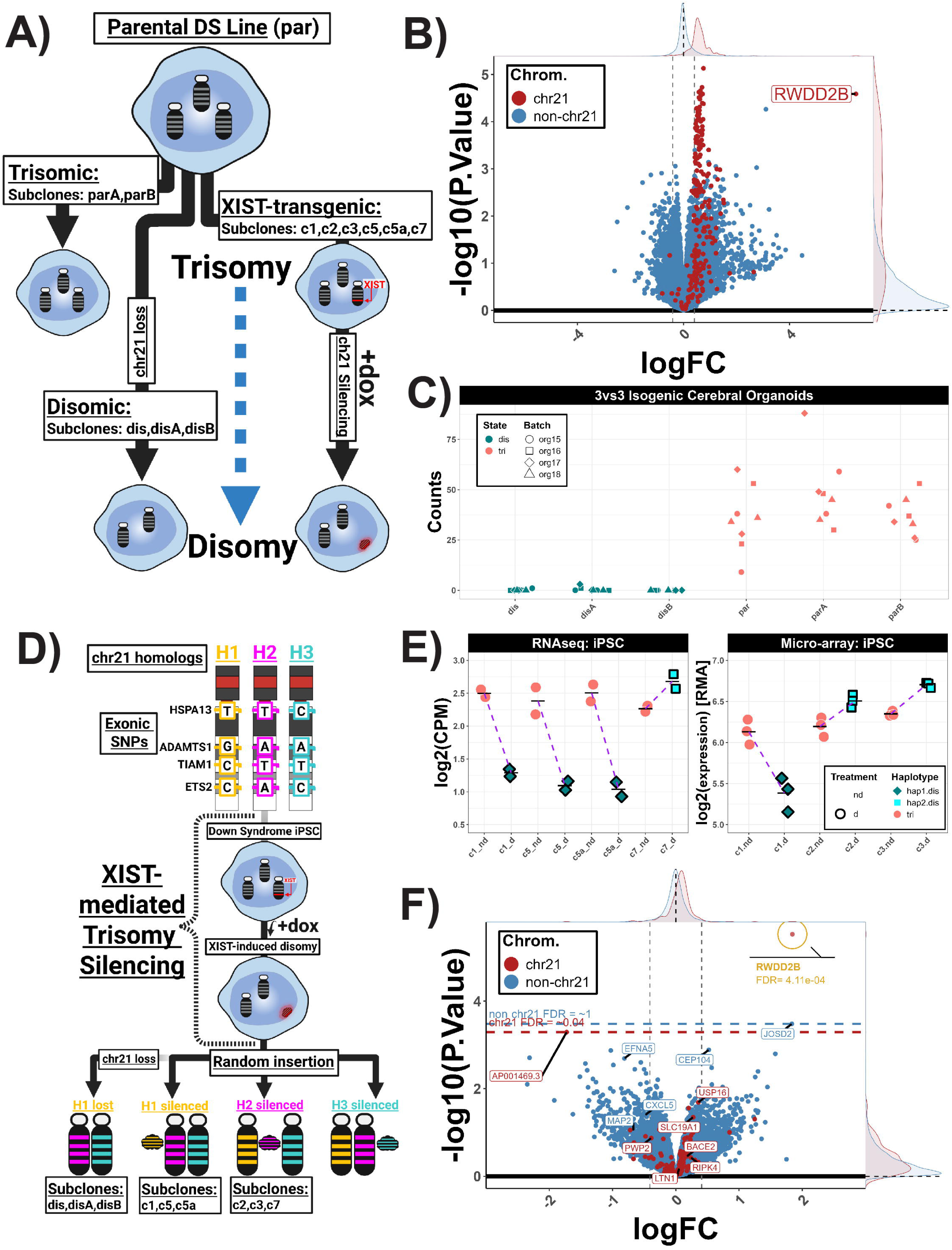
“Selective homolog silencing” links *RWDD2B* expression to specific chr21 homolog. **A)** Isogenic iPSC panel of trisomic, disomic, and XIST-transgenic subclones derived from a single DS individual. **B)** Comparison of three trisomic and three disomic subclones across >1200 organoids showing ∼1.5-fold change in expression of typical chr21 genes, with the notable exception of *RWDD2B*. **C)** Read counts for *RWDD2B* suggested extremely low, potentially off, expression across all disomic samples. **D)** Schematic of “selective homolog silencing”, detailing our clone panel in which specific chr21 homologs are silenced (or lost). **E)** *RWDD2B* expression changes in iPSCs upon chr21 silencing vary based on which particular chr21 homolog is silenced; H1 (c1, c5, c5a) or H2 (c2, c3, c7). **F)** Differential expression analysis of the impact of silencing H1^RWD^ (c1, c5, c5a) vs H2 (c7) in iPSCs shows strong significance for only *RWDD2B*. Volcano plots: Red = chr21, Blue = non-chr21. Density for chr21 and non-chr21 displayed on the axis edges. Vertical lines = +/- log2(1.5-fold change). Horizontal lines = Lowest non-chr21 FDR & lowest non-*RWDD2B* chr21 FDR. Count plots: Pink = Trisomic (or XIST non-induced), Blue = Disomic (or XIST induced), Shape = organoid batch (C) or chr21 haplotype (E), Outlined = Dox treated. Overview diagrams (A+D) made using BioRender.

### Large *RWDD2B* expression changes are not linked to T21 itself but to silencing or loss of a specific chr21 homolog

In multiple transcriptomic datasets from our lab, *RWDD2B* showed exceptionally large fold changes, as seen in a recent study of cerebral organoids comparing our three trisomic and three disomic isogenic sublones^53^ (Fig 1B) and with induced “trisomy silencing” in studies using multiple *XIST*-transgenic iPSCs subclones and derived endothelial cells^41^ (Sup Fig 1A+B). Interestingly, in our data *RWDD2B* was confidently detected at modest levels in our trisomic samples but was barely detectable in the disomic samples (which had spontaneously lost one chr21), causing an outlier fold-change (Fig 1C). We initially thought the large fold-change in *RWDD2B* mRNA might be caused by trisomy itself, such as through amplification caused by higher dosage of chr21 transcription factors (TFs) that positively regulate *RWDD2B*. However most transcription factor binding sites are not clearly defined and thus no clear candidate could be identified^54,55^

The possibility that this marked difference in expression level was due to genetic differences between the three chr21 homologs seemed initially remote, because this was seen in an average of several independent clones, across multiple experiments. However, to address this directly we determined which chr21 homolog was silenced in each of our *XIST*-transgenic clones, as in Jiang et al.^38^, and then re-analyzed data based on the silencing of different chr21 homologs (Fig 1D). While *RWDD2B* was expressed in all transgenic clones to start, after silencing was induced *RWDD2B* mRNA levels dropped sharply only in clones (c1, c5, c5a) that silence the same specific chr21 homolog, termed H1^RWD^. In contrast, one clone (c7) that silences a different chr21 homolog (termed H2) showed little if any reduction in *RWDD2B* post-silencing (Fig 1E; Right). By excluding the *XIST*-transgenic clone targeting H2 from our previous analysis in iPSCs, the fold-change and significance of *RWDD2B* notably increased, with little to no impact on other chr21 genes (data not shown). Similarly, when we analyzed our data to test for the difference between silencing H1^RWD^ or H2 (see Methods), *RWDD2B* was one of just two significant genes; even neighboring genes *LTN1* and *USP16* were not significant, indicating an effect specific to *RWDD2B* (Fig 1F + Sup 1C, Table S4). Since our current iPSC panel only included one clone that targeted H2, we examined earlier microarray data from other transgenic clones^38^, which again showed a sharp drop with silencing of H1^RWD^ in c1 but not in two additional clones (c2, c3) that silenced H2 (Fig 1E; Right). We also determined that each of our three disomic subclones had spontaneously lost H1^RWD^, explaining our findings of negligible *RWDD2B* expression in disomic cerebral organoids. Hence, data from all ten subclones supports that *RWDD2B* is more highly expressed from the H1^RWD^ homolog than the H2/H3 homologs.

### Chr21 homolog-specific transcriptional silencing of specific *RWDD2B* alleles linked to promoter methylation

As shown in Figure 1C, *RWDD2B* mRNA levels drop to extremely low levels with the loss of H1^RWD^, suggesting that *RWDD2B* transcription may be epigenetically silenced in an allele-specific manner. RNA-seq is not well-suited to distinguish transcriptional absence from very low mRNA levels and also will reflect mRNA stability. However, RNA FISH to transcription foci is a well-established approach to examine allele-specific silencing, as used to study X-chromosome inactivation^56–59^. Hence, we examined *RWDD2B* transcription foci in *XIST*-transgenic clones that target different homologs for silencing. In many other studies visualizing transcription foci for various genes, we generally observe RNA foci emanating from all alleles^38,40,59^. While the efficiency of a small probe to detect the *RWDD2B* foci is likely lower than for many genes we have examined (see Methods), only one transcription focus was detected in the majority of *RWDD2B*-expressing trisomic cells, and roughly half of the cells had at least one *RWDD2B* focus. Most importantly, silencing of H1^RWD^ (c5) sharply diminished the number of cells with transcription foci, whereas silencing H2 (c7) had no effect (Fig 2A+B). This was confirmed in two experiments (see Methods), one comparing cultures with or without doxycycline (dox)-induction of *XIST*-based silencing (Fig 2A; Left + Sup Fig 2A) and another comparing *XIST* expressing and non-expressing cells within the same culture, thus controlling for technical variation between slides (e.g. hybridization efficiency) and the potentially influence of dox (Fig 2A; Right). Reinforcing that H1^RWD^ was the expressing homolog, *RWDD2B* foci were essentially absent in two disomic subclones that lost H1^RWD^ (Fig S2B-D).

**Figure 2:**
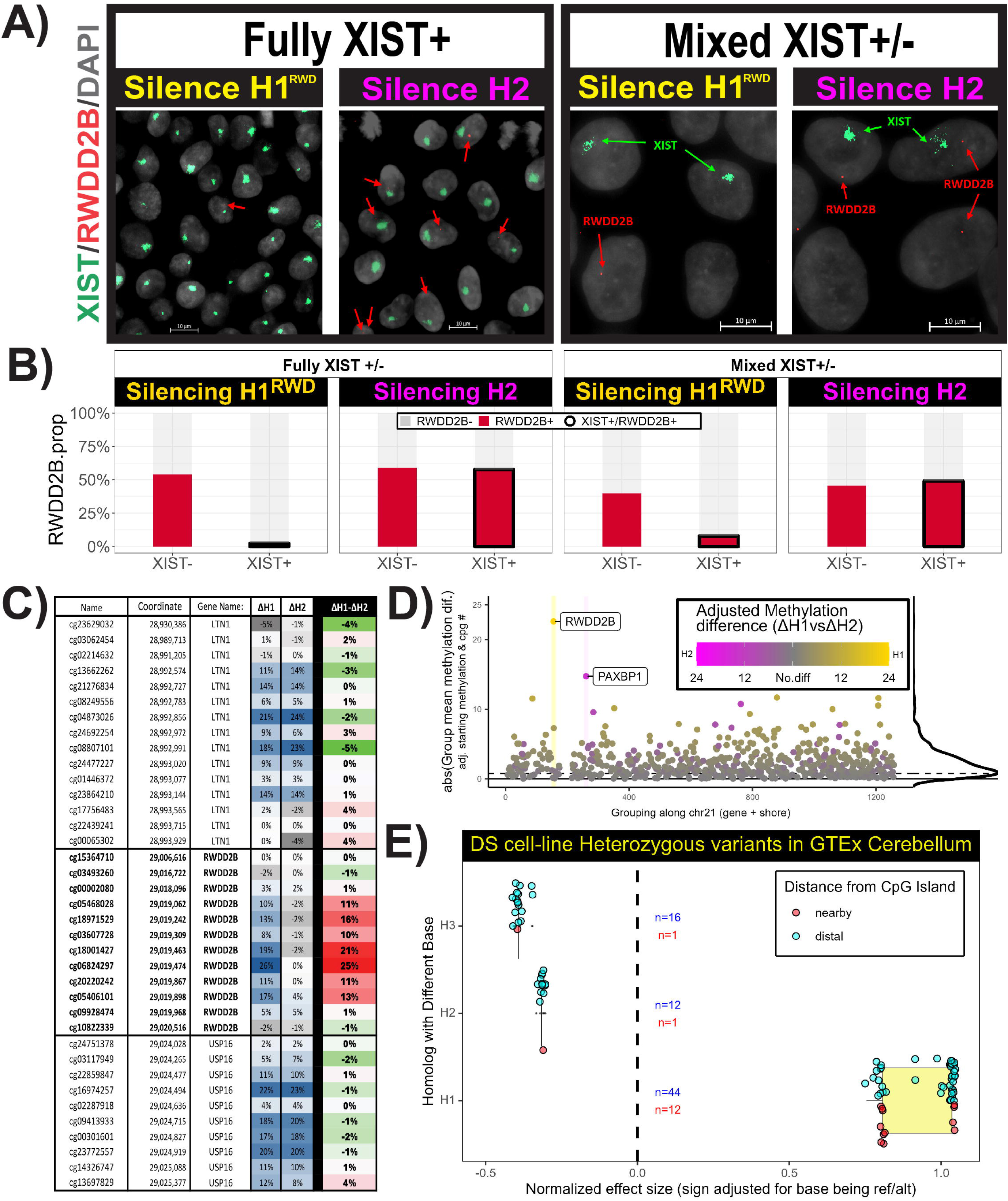
R***W***DD2B **allele-specific transcription linked to difference in CpG methylation and genetic polymorphisms. A)** RNA FISH for *RWDD2B* (Red) and *XIST* (Green) in cultures fully induced for *XIST* and mixed cultures where cells either express *XIST* and silence H1^RWD^ (c5; left) or H2 (c7; right) or did not express *XIST*, remaining functionally trisomic. **B)** RNA FISH quantification in +/- dox and mixed *XIST*-expressing cultures shows *RWDD2B* foci loss only with H1^RWD^ silencing (n >1000 cells per condition). **C)** CpG methylation differences for *RWDD2B* and neighboring genes (*USP16*, *LTN1*) following *XIST*-induced silencing of either H1^RWD^ (c1) or H2 (c2). Blue = methylation increase, Grey = methylation decrease, Green = H1^RWD^ methylation increased less than H2, Red = H1^RWD^ methylation increased more than H2. **D)** Average CpG Island methylation difference across chr21 genes after silencing different chr21 homologs strongly detects difference for *RWDD2B*. *PAXBP1* was the only other strong outlier. Yellow = stronger methylation difference when silencing H1^RWD^, Pink = stronger methylation difference when silencing H2. E) Phased polymorphisms heterozygous in our cell-line in a 100kb region flanking *RWDD2B* and with variants different on H1^RWD^ from H2/H3 strongly correlate with stronger, positive, normalized effect sizes in GTEx cerebellum with H2/H3 correlating with weaker, negative, effect sizes. The narrowed most likely candidates are within 5.5Kb of the *RWDD2B* CpG island/Promoter Region (red).

We note that while two trisomic subclones examined by FISH showed primarily only a single transcription focus, a small percentage of cells had two, and ∼1% of cells had three (Fig S2B-D). This suggests there may be some epigenetic instability that affects the transcriptional silencing of particular alleles. Nonetheless, results indicate *RWDD2B* transcription is essentially monoallelic in our iPSCs and their derivatives, with expression being linked to the H1^RWD^ homolog.

We next considered that the *RWDD2B* alleles on H2 and H3 may be epigenetically silenced by increased promoter CpG island methylation. To investigate this, we capitalized on the fact, established in studies of X-inactivation, that *XIST*-based silencing increases CpG methylation in gene promoters^28,29,56^. If the *RWDD2B* promoter on H2 and H3 were already strongly methylated and silenced, then *XIST*-mediated silencing may increase methylation only on H1^RWD^, where methylation may be lower to start with. For this we re-examined the *RWDD2B* locus in DNA methylation array data from our earlier study which had shown that *XIST*-mediated chr21 silencing increased promoter methylation for almost all of 143 well-expressed chr21 genes in two transgenic clones (Table S5)^38^. Interestingly, at that time the data for *RWDD2B* was considered uninterpretable because the two transgenic clones gave conflicting results. However, this difference can now be explained by differences between the homologs: when silencing H1^RWD^ (c1), *RWDD2B* promoter methylation increased, consistent with H1^RWD^ carrying the transcriptionally active allele (Fig 2C). In contrast, methylation did not change when silencing H2 (c2) as it carries an already transcriptionally silent allele, consistent with RNA-seq and FISH. Importantly, almost all other chr21 genes (including neighboring genes *LTN1* and *USP16*) showed similarly increased methylation upon *XIST*-induction irrespective of which homolog was silenced, further underscoring the unusual nature of *RWDD2B* within our cells (Fig 2D). Collectively, results show that selective homolog silencing enabled the discovery of distinct epigenetic regulation of CpG methylation and transcriptional status between *RWDD2B* alleles.

### Polymorphisms in/near *RWDD2B* promoter on H1^RWD^ versus H2/H3 strongly correlate with higher expression

We next considered whether this epigenetic difference in *RWDD2B* reflects underlying genetic differences, and thus we determined the local haplotypes of H1^RWD^ versus H2 and H3 in our cells and examined differences with respect to eQTLs quantified in GTEx (see Methods). As this strong impact on expression was limited to *RWDD2B*, not strongly impacting even neighboring genes (Fig S1C), we focused on heterozygous polymorphisms within a 50kb region flanking *RWDD2B*. Phased whole genome sequencing for this region in our trisomic line (see Methods) demonstrated that polymorphisms different on H1^RWD^ compared to H2 and H3 show markedly stronger effect size in GTEx as compared to those shared between H1^RWD^ and either H2 or H3. This is shown in the cerebellum, the tissue with the strongest effect sizes in GTEx (Fig 2E). The effect directionality between the homologs further reinforces that this finding is genetically driven, as eQTLs different on H1^RWD^ correlate with higher *RWDD2B* expression whereas those different on either H2 or H3 correlate with lower expression. Thus, the difference in transcription from *RWDD2B* alleles is associated with genetic polymorphisms, not simply explained by gene imprinting during gametogenesis or random monoallelic expression (both of which are not generally controlled by polymorphisms)^28,60–62^. Interestingly, epigenetic instability of genes that show variable escape (or not) from X-chromosome silencing has also been suggested to be influenced by genetic polymorphisms^63^.

This approach using genomic analyses on cells from a single individual also enabled us to narrow which *RWDD2B* eQTLs in GTEx likely relate to this large expression difference, distinct from those that more modestly modulate expression. Furthermore, our methylation data indicates that only the CpGs in the center of the *RWDD2B* regulatory region (promoter + CpG Island) showed differences, thus the responsible variants are most likely in or proximal to this region^64^. By narrowing down to variants specific to H1^RWD^ and those proximal to this regulatory/methylation region (Fig 2E; Red vs Blue) this approach can reduce the >1,000 significant *RWDD2B* eQTLs in GTEx to the 68 different on H1^RWD^, of which just 13 (1 not in GTEx) were within 5.5Kb of *RWDD2B*’s regulatory region and likely responsible for altered methylation and thereby expression (i.e. meQTL) (Table S6 + Fig S3B). We note 18 heterozygous polymorphisms were not phased in our cell line and were not tested in GTEx V10. This demonstrates how our cell-based approach from a single individual can complement large studies across many individuals to help narrow the most likely candidates for relevant phenotypes.

### RWDD2B regulation by meQTLs is cell-type specific with strongest effects in brain

Results above discovered that genetic variants can strongly influence *RWDD2B* transcriptional status, as evidenced in both iPSCs and cerebral organoids from our cell-line. Based on GTEx data from various tissues, variants in our cell-line that were different on H1^RWD^ have the strongest effect size in brain tissues and weakest in skeletal muscle (Fig 3A). GTEx expression data is primarily from human tissues, which contain multiple cell-types, whereas the iPSC approach used here enables examination of eQTL effects in specific cell types. For example, by examining endothelial cells differentiated from the same iPSC subclones (Fig 3B, Table S7) we find the same strong effect on *RWDD2B* expression (when comparing the silencing of H1^RWD^ vs H2), which could not be determined from GTEx.

**Figure 3:**
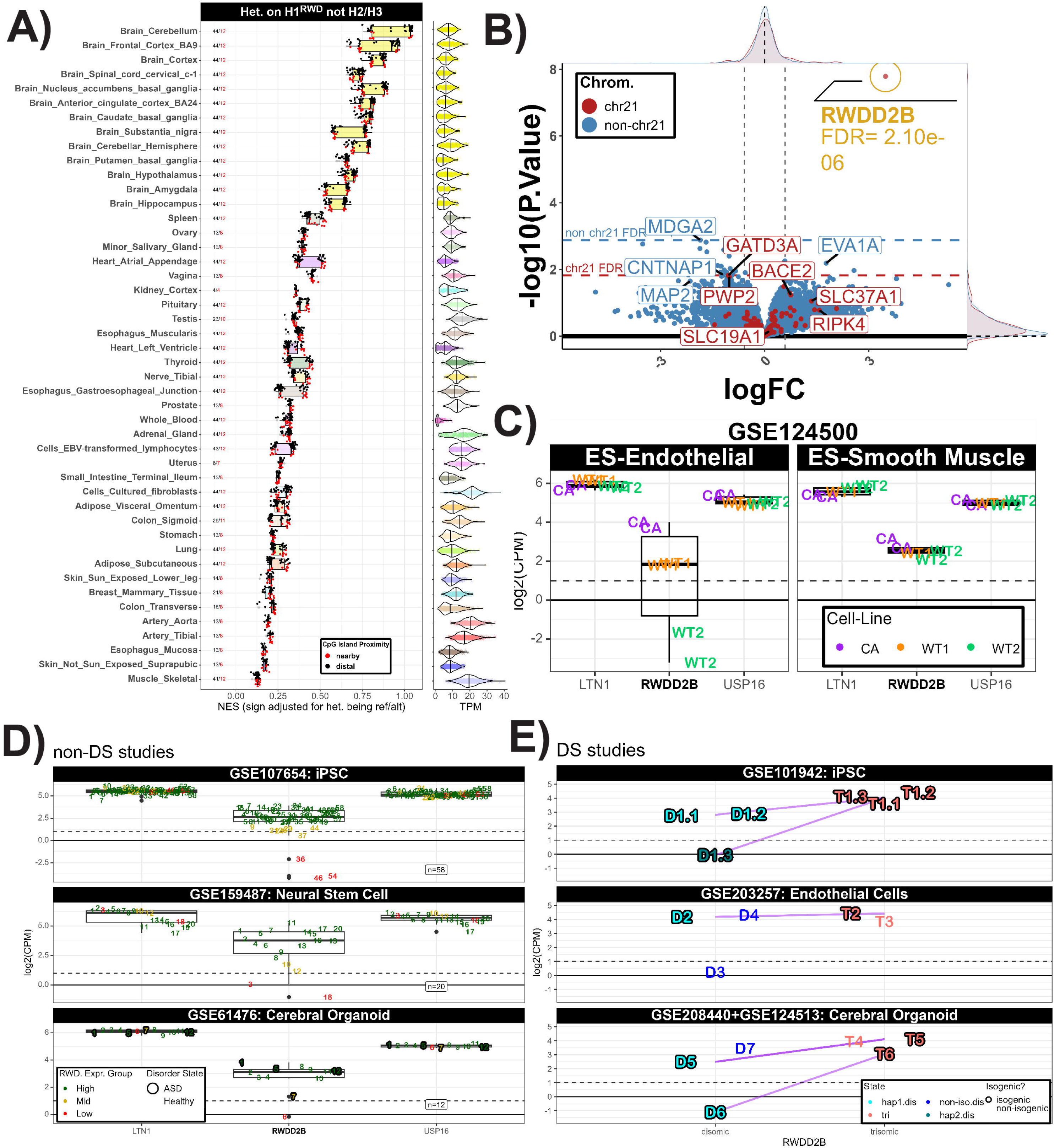
Effects of genetic polymorphisms on *RWDD2B* expression are cell-type specific and expression variation is seen across many datasets. A) Focusing only on eQTLs from GTEx for which the variant at H1^RWD^ differs from H2/H3, normalized effect sizes of eQTLs are largest in neural tissue (yellow) and smallest skeletal muscle, with some tendency for tissues with larger effect size to have lower average *RWDD2B* expression (TPM shown on right). Red = promoter proximal, Black = promoter distal. **B)** Differential expression analysis for the impact of silencing H1^RWD^ vs H2 in iPSC-derived endothelial cells also shows that *RWDD2B* expression is from H1^RWD^. C) *RWDD2B* expression data in 3 non-isogenic hES cell lines show large differences between individual when differentiated to endothelial cells, but not when differentiated to smooth muscle, with little variation for neighboring genes. D) *RWDD2B* expression data from three different studies, using cell-lines representing ∼90 euploid individuals, each show a similar pattern of High (Green), Mid (Yellow), and Low (Red) expression not seen for neighboring genes. Outlined = Autism spectrum disorder (ASD). E) *RWDD2B* expression data from four different studies comparing trisomic and non-trisomic iPSCs or derived cell-types show that the loss of a homolog can result a range in expression changes from no change to a sharp drop. For cerebral organoid data, T5/D5 come from GSE208440, all others from GSE124513. In top panel, isogenic pairs (outlined) that lost different homologs are colored light vs dark blue (clones which lost the RWD associated homolog are dark blue. Purple lines connect the average expression of isogenic trisomic-disomic pairs for the each chr21 homolog when applicable.

Collectively, the above findings indicate that the “functional dosage” for *RWDD2B* is less tightly predicted by the gene copy number compared to most genes, which has implications for DS and the general population. Evidence that *RWDD2B* expression levels can have phenotypic consequences comes from studies reporting that meQTLs associated with reduced *RWDD2B* expression levels are also associated with higher risk for osteoarthritis (OA), first reported in Parker et al.^26^ and then affirmed by others^44,45^. Parker et al.^26^ reported more modest differences in *RWDD2B* mRNA levels than we observe in our data; however, this was based on averages of tissues (which contain multiple cell types) mostly from a large population, whereas the approach here made it possible to examine expression across cells of a pure cell type and even examine individual alleles. Thus, allowing us to more completely assess the allele-specific effect size. As suggested by GTEx data, samples such as whole blood may not display substantial eQTL effect sizes for *RWDD2B* (Fig 3A + Fig S3A). Consistent with this, in whole-blood RNA-seq data^65^ from the Include Project (https://portal.includedcc.org/), we observed no large deviations in *RWDD2B* expression levels among individuals either with or without DS (Fig S4A). Results presented here also indicate polymorphisms can confer greater susceptibility to epigenetic silencing; this may be important for understanding environment-gene interactions in disease (see Discussion).

These findings suggest the important possibility that in some individuals *RWDD2B* expression may be very low in certain cell types, including those that comprise the neural tissue in cortical brain organoids^66,67^ (Fig 1B+C) (See Discussion). Given that this could have broader implications, we asked whether marked differences in *RWDD2B* expression could be seen in cell lines from other individuals (both euploid and trisomic)^66–70^. DNA sequencing data was not available for almost all studies examined, but RNA-seq results indicate strong differences in *RWDD2B* levels consistent with our findings of allele-specific differences. For example, in three non-isogenic embryonic stem cell (ESC) lines (euploid) differentiated into endothelial cells^69^, we found marked differences in *RWDD2B* RNA levels (Fig 3C), consistent with our results indicating strong eQTLs in endothelial cells. Importantly, the same three non-isogenic cell lines showed no difference in *RWDD2B* levels when differentiated into heart muscle, as would be predicted from the much weaker effects of the GTEx eQTLs in heart muscle (Fig 3C). Further, in datasets from cells representing many euploid individuals, from iPSCs, iPSC-derived neural progenitors and cortical organoids^66–68^, we found that a subset of cell samples (from different individuals) have markedly lower *RWDD2B* expression than most others (Fig 3D); there was also some suggestion of an intermediate level (See Methods). In contrast, this marked variation in RNA levels was not seen for the two neighboring genes (*LTN1* and *USP16*), which showed more tightly clustered mRNA levels. As each gene was compared from the same samples and RNA-seq analysis across various studies, the greater differences in *RWDD2B* are not technical artifacts, but are consistent with evidence above of strong *RWDD2B* meQTLs that can make some alleles more prone to epigenetic silencing (as shown in our cells by RNA FISH). We also found large *RWDD2B* expression differences in a study of ESCs from many individuals (Fig S4B) (See Discussion)^70^.

Further evidence comes from other DS studies comparing isogenic trisomic versus derived isogenic disomic lines (which lost a chr21). In several datasets, we observed a similar marked drop in *RWDD2B* expression linked to loss of a chr21 homolog. As summarized in Supplement Table S8, our analysis of all available isogenic trisomic/disomic pairs in GEO supports that *RWDD2B* expression level commonly does not simply reflect copy number^71–80^. This data supports that there could be three possible outcomes with loss of a chr21 homolog in T21; an expected decrease in *RWDD2B* expression (loss of a highly expressed allele with >1 highly expressed alleles remaining), no significant decrease (loss of an essentially non-expressing allele), or a marked decrease (loss of the only highly expressed allele), as observed in our cell line. Examples are provided in Figure 3E. In the case of data from Gonzales et al.^73^, we could affirm that loss of a specific homolog was linked to a stark drop in *RWDD2B* expression, whereas loss of a different homolog was not (Fig 3E; Top). Our examination of data from the Pinter lab’s XIST-based silencing of one particular chr21 homolog^81^ also found that this resulted in a stark, outlier, drop in *RWDD2B* expression (data not shown).

### Broader methodological potential of “selective homolog silencing” to study of polymorphisms

Our findings regarding *RWDD2B* demonstrate a new concept and experimental strategy, “selective homolog silencing”, which we suggest could be a valuable complement to large population-based studies. Most eQTLs have more modest effects than the large-effect size of *RWDD2B*, which was evident from analysis of just a few clones (from one individual). However, this approach could be expanded to 1) identify more modest chr21 eQTLs, and 2) study the *trans* effects of chr21 polymorphisms in different cell types (Fig 4A+ Fig S4C). To illustrate these potential applications, we show examples of potential candidates for chr21 genes with *cis* eQTLs, or off-chr21 genes impacted in *trans* by chr21 polymorphisms. These tentative candidates are based on limited data (designed for another purpose), but the experimental power could be improved by including both more clones (targeting specific homologs) and cells from several individuals. Interestingly, our analysis for “selective homolog loss” (see Methods) using data from Gonzales et al.^73^ revealed *RWDD2B* as the only chr21 gene whose expression level differed significantly depending on which copy of chr21 was lost (Fig 4B), highlighting its unusual nature. Finally, this approach could also be potentially extended to other chromosomes or regions of interest (see Discussion).

**Figure 4:**
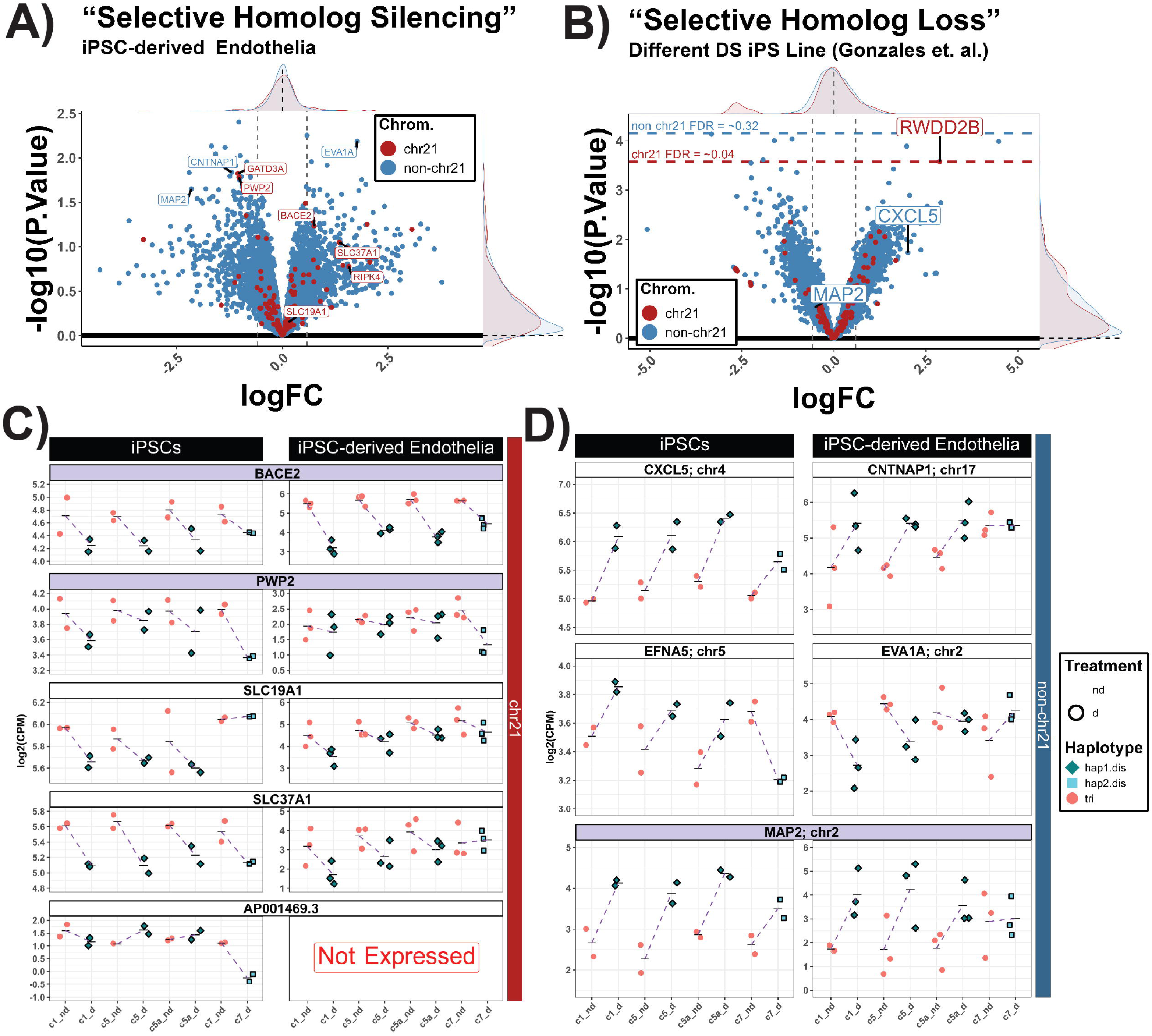
B**r**oader **potential use of selective homology silencing or loss to study eQTLs. A**) Volcano plot comparing silencing of H1^RWD^ vs H2 to identify candidate chr21 influenced by polymorphisms on chr21 QTLs with more modest effects than RWDD2B (RWDD2B and non-chr21 genes with log2(CPM) < 2 removed) in iPSC- derived endothelia. **B)** Volcano plot for “selective homolog loss” using iPSC data from Link et al. (non-chr21 genes with log2(CPM) < 2 removed) strongly indicating difference for *RWDD2B*. **C)** Count plots for several candidates chr21 eQTLs in iPSCs and derived endothelial cells. <u>Top</u>) BACE2 and PWP2 are two genes with consistency in both cell types. <u>Middle</u>) The two SLC genes are examples that appear to have cell-type specific effects. <u>Bottom</u>) AP001469.3 was the only other significant chr21 gene identified. **D)** Count plots for preliminary genome-wide candidates whose expression corresponded with selective homolog silencing; with MAP2 showing consistency in both cell types (bottom). Volcano plots: Red = chr21, Blue = non-chr21, Density for chr21 and non-chr21 displayed on the axis edges. For B) Horizontal lines = Lowest non-chr21 FDR & lowest chr21 FDR. Count plots: Pink = Trisomic (or XIST non-induced), Blue = Disomic (or XIST induced), Shape = chr21 haplotype (iPSC), Outlined = Dox treated. Purple title = consistent between cell types.

Among the chr21 candidates with expression differences between H1^RWD^ & H2 in our cell line are *PWP2* and *RIPK4,* which are both among the chr21 genes identified to have potential eQTLs in iPSCs across ∼220 lines in DeBoever et al.^33^ Interestingly, *RWDD2B* was among the top 5 chr21 eQTLs^33^. Another chr21 candidate we identified, *AP0001469.3*, was the only other significant gene identified, which acted inversely to *RWDD2B* (Fig 4C; Bottom). Interestingly, in both endothelial cells and iPSCs, eQTLs were implicated for *BACE2*, *PWP2*, and *RIPK4*, whereas *SLC19A1* and *SLC37A1* eQTLs appeared cell-type specific (i.e only showing differences in one cell-type) (Fig 4C). Since *RWDD2B* and *USP16* share back-to-back promoters, *RWDD2B*’s regulatory region might affect *USP16*—as suggested by our iPSC data—but endothelial data showed no difference, so further testing is needed (Fig S1C+ Fig S3B).

Ultimately the challenge is to understand the biological connection between specific polymorphisms and clinical phenotypes. To understand how the cellular effects of an eQTL can connect to the clinical phenotype, we illustrate how this strategy can generate hypotheses to help fill the knowledge gap. Hence, we looked genome-wide for non-chr21 genes whose expression change differed between silencing of H1 and H2, and identified examples in both iPSCs and endothelial cells (Fig 4D). Interestingly, *MAP2* was a top candidate in endothelial cells with a consistent, but mild, change in iPSCs (Fig 4D; Bottom) and MAP2 has been linked to dendritic pathology in DS^82,83^.

While this strategy can suggest non-chr21 gene whose expression levels are affected in *trans* by variants on chr21, it does not directly reveal specific SNPs or genes on chr21 that are responsible for this. However, in datasets for which *RWDD2B* is by far the chr21 gene with the strongest expression difference between chr21 homologs, it is plausible that some expression changes in non-chr21 genes may be due to levels of *RWDD2B* itself, which could potentially give insight into why meQTLs for *RWDD2B* are associated with OA^26,45,46^. For example, *CXCL5* was a candidate (top 200 non-chr21 genes by P-Value) for *trans* chr21 effects in our iPSC data (Fig 4C + Fig S4C) and was suggestive in our “selective homolog loss” analysis of data from Gonzales et al.^73^ with stronger differences from T21 subclones when losing high *RWDD2B* expressing chr21 homolog (Fig 4B + Fig S4D). Intriguingly, CXCL5 is strongly involved in joint inflammation in arthritis, including OA^84–86^, and has been noted to as a therapeutic target in arthritis^87^. Another preliminary example from our data, *EFNA5*, a member of the ephrin family, is strongly expressed in the syncytial joint, and ephrin signaling dysregulation may be involved in joint disease pathology and be implicated in inflammation^88–91^. These preliminary examples illustrate how this strategy could be extended to gain insight into how polymorphisms give rise to relevant cellular phenotypes, which could be extended more broadly (see Discussion).

## Discussion

This study has significance for both biological findings regarding RWDD2B regulation and the demonstration of “selective homolog silencing” as a novel approach to identify eQTLs and investigate cell-type specific effects. By manipulating which chr21 homologs are silenced by *XIST* RNA we discovered the on-off transcriptional state of *RWDD2B* alleles can be epigenetically regulated by polymorphic variation based on cells from a single individual. In essence, functional dosage of *RWDD2B* was impacted by allelic silencing in the three cell-types we examined: iPSCs, cerebral organoids, and endothelial cells^41,53^. Variation in *RWDD2B* expression is relevant to the general population, as evidenced by the link with osteoartritis^26,45,46^, but the range in expression levels can be even wider in the DS population (from 0-3 expressing alleles). It is notable that GTEx eQTLs for *RWDD2B* have largest effect sizes in the brain, consistent with the large effect size seen in our neural organoids, raising the possibility that some individuals (euploid or trisomic) may express little (or no) RWDD2B in many neural cells of the developing brain. Even if not essential for these cell-types, differences in expression might still affect brain function, development, or health in some way. However, for reasons explained below, we suggest the hypothesis that RWDD2B may play a role in immune and inflammatory responses.

Human genomes are replete with genetic variation that often has quite subtle effects on gene expression levels, which in turn may not have significant biological impact. However, *RWDD2B* appears to be unusual in that a relatively common polymorphism(s) can influence the on-off state of transcription in various cell-types. While the functions of RWDD2B are unknown, several observations support that it is likely dosage sensitive. Most importantly, modestly decreased *RWDD2B* expression linked to meQTLs is a risk factor for osteoarthritis (OA)^26,45,46^. On the other hand, *RWDD2B* was among 11 of 163 chr21 genes tested that consistently produced a phenotype when overexpressed in zebrafish, impacting somite morphology^42,43^. Furthermore, this difference in *RWDD2B* expression appears to be cell type specific, which may suggest that the functional dosage of RWDD2B “matters”, at least for the development and/or function of certain tissues. Finally, a recent study discovered that RWDD2B protein levels are highly regulated, as RWDD2B was unexpectedly identified as the top target for degradation by TRIM7, a ubiquitin ligase that regulates turn-over of factors involved in the immune response^44^.

While RWDD2B is not well studied, the few reported biological interactions we found implicate it in immune-related functions. TRIM7 (as described above) and the TRIM protein family are key regulators of the viral immune response and thus are heavily studied in relation to autoimmunity and inflammation^92–94^. Epigenetic regulation of *RWDD2B* was also reported to be altered following the humoral immune response^95^. In addition, RWDD2B was shown to interact with GADD45G^96–99^, a negative regulator of the JAK-STAT signaling pathway which is centrally involved in regulation of immune functions^100^. Most individuals with DS have high levels of inflammatory markers in their blood, which is thought to be linked to triplication of the four interferon receptor genes on chr21^7,9,10^. Chronic inflammation is now thought to be a major driver of OA in the general population^8,47,48^. Hence, RWDD2B levels may modify the risks of OA and other immune-related conditions in the general population, but particularly so in DS where inflammation, viral susceptibility, and autoimmunity are frequent and major clinical challenges that vary between individuals. Interestingly, OA of the hip joint is a DS co-morbidity^1,4,5^ suggesting consideration of how differences in RWDD2B levels could exacerbate or mitigate the variable impacts of T21, including OA and hyperactive immune response. TRIM7 also has broader functions, and is suggested to promote heart muscle proliferation and migration, potentially relevant to congenital heart defects in DS^101^. Similarly, GADD45 and the JAK-STAT pathway both relate to cellular senescence, which is known to be increased in DS^99,100,102,103^, and JAK-STAT itself has been reported to be disrupted in DS^104,105106^. While these connections to RWDD2B are speculative at this point, we believe these findings elevate the potential relevance of RWDD2B in DS.

Very few studies focus on RWDD2B, but we found RWDD2B listed (among many genes) or mentioned in a few epigenetic studies of Alzheimer’s disease (AD), humoral immune response to influenza vaccination, breast cancer, and insomnia^95,107–110^. Other evidence shows vorinostat (SASHA) treatment in T-cells strikingly increased *RWDD2B* expression 11-fold^111^. While commonly used as an HDAC inhibitor, SASHA also impacts methyltransferase and can trigger DNA demethylation^112–115^. *RWDD2B* polymorphisms were initially identified in a search for mQTLs^116^ and subsequently meQTLs^26,45,46^ associated with OA and, consistent with this, our findings suggest these polymorphisms influence susceptibility to epigenetic silencing. Our analysis of allele-specific transcription in single cells from a single individual provides evidence that *RWDD2B* levels are not simply modulated by genetic factors or other aspects such as RNA turnover, but that genetic variants alters promoter CpG methylation resulting in allele-specific silencing of *RWDD2B*. This “on-off” regulation is linked to polymorphic sites (with quantified effect sizes in GTEx) different on particular homologs in our cell line and thus is distinct from parental imprinting or random mono-allelic expression. RWDD2B has not been flagged in several studies that looked for imprinted chr21 genes^117–119^ and was reported to be bi-allelic in a search for mono-allelically expressed genes^120^. In searching the literature, we found extremely limited reports where altered methylation was described to confer “on-off” transcriptional status associated with polymorphisms, as we observed for *RWDD2B*. Moreover, these were thought to impact regional methylation which impacted transcription factor binding^60,62,121–126^. We also note that this difference for *RWDD2B* may not relate to a single variant but rather may be due to the co-occurrence of several variants, which is difficult to examine, though methods are emerging^34,35^. While polymorphism-driven propensity for epigenetic transcriptional silencing is likely not common, it is also the case that it was made more apparent by the cell-based approach used here, as discussed below.

It is interesting to note that the difference between *RWDD2B* alleles in terms of DNA methylation (and corresponding transcription status) was evident in pluripotent cells, given that reprogramming of somatic cells to a pluripotent state generally involves removal of DNA methylation^127,128^. However, there can be slight differences between the full extent of epigenomic reprogramming for each iPSC clone, with some newer procedures reported to produce more “naïve” iPS cells^129,130^. The ten iPSC subclones used in this study were isolated from a trisomic iPSC line about ten years ago^38^, and all ten lines are consistent with *RWDD2B* transcription primarily from H1^RWD^ and very little (if any) from H2 and H3. Collectively, data supports that genetic polymorphisms confer greater propensity for H2 and H3 alleles to become methylated and/or H1^RWD^ to resist methylation. We also provide evidence for allelic differences in *RWDD2B* expression in ES-derived lines (Fig 3C) and in a larger pool of ES lines from several independent individuals, which were not generated by reprogramming (Sup Fig 4B).

The approach we describe here, to manipulate expression of a whole chromosome, is limited in resolution and only informative for polymorphisms heterozygous in each individual’s cells. However, when coupled with GTEx data or potentially other methods for allele-specific expression analysis^60^, even results from just one iPSC line can expand what can be investigated in several ways. First, GTEx data is based on samples from a large population from tissues with multiple cell-types, whereas the approach here allows analysis of single cells and even single alleles, which can then be correlated with GTEx data. We show that primarily one *RWDD2B* allele was transcribed, while the other alleles were transcribed at much lower frequency, if at all. Low levels in RNA-seq would not provide the same information as evaluation of transcription foci from individual cells. Second, cell-based analysis from a single individual can assess cell-type specificity in more pure cell populations after iPSC differentiation than analysis of tissues, which has required hundreds of iPSC-derived lines in other studies, limiting practical extensions^30–33^. As illustrated by *RWDD2B*, the effects of eQTLs can be highly cell-type specific, so effects can be diluted or even missed in tissues. For example, our results suggest larger effect sizes for *RWDD2B* than those seen in Parker et al.^26^, which was based on RNA levels in heterogenous tissues from many people. Third, by identifying heterozygous SNPs specific to each homolog in our cells, this greatly narrowed the likely candidates impacting *RWDD2B* expression from a much larger pool based on GTEx data alone, and identified a handful of candidates not tested in GTEx or identified as eQTLs in other studies^26,33^. Fourth, focusing on just the narrowed candidate SNPs, we see a larger effect size (e.g. in the brain) than estimated from just all significant *RWDD2B* eQTLs in GTEx, which reflects an average of many SNPs, including those weaker impacts. Our analysis also provides information regarding the co-occurrence of SNPs as a haplotype.

This framework to study polymorphisms could be expanded to study other chromosomes or smaller chromosome regions. Not only could this be extended to other viable trisomies (e.g. 13 and 18), silencing of non-trisomic autosomes is likely also possible, despite whole chromosome monosomy having longer-term viability effects on cells in culture. However, our lab recently developed an *XIST*-based approach that represses only a small, targeted, region of a chromosome (Valledor et. al.^59^ and unpublished), which could enable “localized haplotype silencing”, allowing for *cis* and *trans* effects to be studied for specific regions. Whatever the approach, we believe effects of polymorphisms in cells are best studied using an inducible approach, to minimize other sources of noise that can arise even between isogenic lines (i.e. epigenetic and genetic drift^131–133)^.

Perhaps most importantly, a mechanism to manipulate homolog/haplotype expression in cells or organoid tissues provides a useful approach for hypothesis generation to help bridge clinical phenotypes (implicated by GWAS) to the underlying cell biology^36,37^. For example, our limited data suggests the possibility of trans-effects on *CXCL5* and *EFNA5*, both of which could connect to immune function and/or OA. Another example of a chr21 with modest eQTLs, *BACE2*, was suggested in our data, and studies have suggested increased *BACE2* may reduce penetrance of AD in DS^134^. *BACE2* eQTLs have also been correlated with the age of AD onset^135^ and amyloid levels^136^, and several of these *BACE2* variants were heterozygous in our cell-line. Both examples serve to illustrate how expansion of the approach shown here can generate hypotheses for further study.

## Methods

### DS iPSC panel creation

While fully detailed in prior publications^38^ and briefly summarized in the results, the panel of induced pluripotent stem cells (iPSCs) described here was generated from a single male individual with Down syndrome (DS/T21), reprogrammed from fibroblasts by George Q. Daley^137^, which we subcloned after an initial electroporation event. Hence the *XIST*-transgenic subclones, trisomic subclones (which did not integrate the *XIST* transgene), and disomic subclones (which lost a single chromosome 21 (chr21) homolog and did not integrate the *XIST* transgene) are all isogenic and generated from the same iPSC line and initial integration event. These subclones were grown and expanded for initial frozen stocks, which have been thawed and subsequently expanded over the years as required for experiments.

### iPSC culture and cell fixation

All iPSCs were cultured with Essential 8 medium (ThermoFisher) in a feeder-free condition on vitronectin-coated plates. Cells were maintained at 37°C, 20% O_2_, and 5% CO_2_, and passaged every 3–5 days (∼80% confluency) using 500μM EDTA in in 1x phosphate buffered saline (PBS). To test *RWDD2B* transcription via RNA fluorescence in situ hybridization (FISH), subclones with *XIST* targeting either ch21 homolog H1^RWD^ or H2 (c5 and c7 respectively) were thawed, expanded, and treated with 500ng/mL doxycycline (dox) for two weeks to ensure full chr21 silencing. However, *XIST* RNA expression was found to be inconsistent across the cell population, due to transgene silencing^40^, hence cells were re-selected using puromycin (3μg/mL) (the TET3G transgene selection marker) for one week, producing highly pure *XIST*-expressing cultures. RNA FISH for *XIST* and *RWDD2B* was performed one week after puromycin removal. Selection with and removal of puromycin was conducted while still in the presence or absence of dox. Later, we realized that cultures with mixed *XIST*-expressing cell populations allowed simultaneous analysis of induced and non-induced cells on the same coverslip, minimizing (or eliminating) technical variation; thus, a second RNA FISH experiment for *XIST* and *RWDD2B* was performed with these cultures. Finally, trisomic (parA, parB) and disomic (dis, disB) subclones were thawed, expanded, and used for confirmatory *RWDD2B* RNA FISH.

For each experiment, cells were seeded onto coverslips two days before RNA FISH, then processed using established protocols^38,138^. Briefly, coverslips were coated with attachment protein, seeded with cells, cultured for two days, then extracted with 0.5% Triton X in 10 mM vanadyl ribonuclease complex (VRC) for 3 minutes, fixed with 4% paraformaldehyde (PFA) in 1x PBS for 10 minutes, and stored in cold 70% ethanol until used.

### RWDD2B RNA FISH and XIST/RWDD2B co-detection, imaging, and quantification

*XIST* was detected using a plasmid containing a ∼9kb fragment (introns 5–8) labeled by nick translation with Digoxigenin-dUTP (Roche), producing ∼100 bp probes spanning exonic and intronic regions. *RWDD2B* could not be detected with BAC-derived probes due to those also containing sequence from the flanking (and more highly expressed) genes and cDNA derived probes were also insufficient for detection. Therefore, a custom Stellaris probe (Biosearch Technologies) of ∼20 bp oligos was used. For *RWDD2B* detection, hybridization employed the manufacturer’s buffers and protocols but the RNA hybridization mix was modified to contain 15% formamide and 125ng/μL human COT-1 DNA to block non-specific binding. Optimal *RWDD2B* signal was achieved with a 5-hour hybridization; longer incubations drastically reduced signal-to-noise. For co-detection, *XIST* was first hybridized following our established protocol^138^, fixed with 4% PFA, and then *RWDD2B* was hybridized as described above. DAPI was used to detect DNA. RNA FISH images used a Zeiss AxioObserver 7 with a Flash 4.0 LT CMOS camera (Hamamatsu). Images were maximal image projections (MIPs) made from z-stack images following deconvolution (adjustable, otherwise default) in *Zeiss*. Brightness and contrast were corrected in Zeiss to best represent what was observed by eye (for *XIST* + DAPI) and optimize for *RWDD2B* clarity. Cells were scored from images in Fiji after automatic brightness and color adjustment^139,140^.

### Brief description of previous sequencing studies using this iPSC panel and differentiated derivatives

While full RNA sequencing methods and differentiation protocols are detailed in the relevant publications, a brief overview is provided here. iPSC sequencing was conducted on four *XIST*-transgenic subclones (c1, c5, c5a, c7) and three isogenic trisomic (par, parA, parB) and disomic (dis, disA, disB) subclones after approximately one week of doxycycline induction (and non-induced controls), across two independent replicates^41^. For this experiment, the percentage of cells with *XIST* induction was reduced due to transgene silencing^40^, dampening chr21 fold-changes. Re-selecting for *TET3G* expression, using its selectable marker (puromycin), was used in subsequent iPSC-derived endothelial experiments^41^. Endothelial differentiation followed a 12-day differentiation protocol, which was replicated three times, differentiation variation likely results in higher variability seen count plots. Cortical organoid experiments involved differentiating our isogenic trisomic and disomic subclones for 90 days across four independent batches with two replicates per batch, totaling >1,200 organoids^53^.

For each experiment RNA was extracted using Trizol Reagent (ThermoFisher) with 100% ethanol for precipitation. RNA was treated with DNase I (Roche) and RNasin Plus (Promega), for DNA digestion, and then purified with the RNeasy Mini Kit, following the manufacturer’s protocol. RNA quality was assessed using a Fragment Analyzer. Sequencing libraries were prepared using the NEBNext Ultra II Directional RNA Kit, Poly (A) mRNA Isolation Module, and Multiplex Oligos for Illumina (New England Biolabs). Sequencing was performed by the UMMS Deep Sequencing Core. Libraries were aligned to GRCh38 using HISAT2^141^ and mapped reads were counted using featureCounts^142^, excluding multi-mapped reads.

### Gene expression analysis across RNA-seq datasets

For RNA sequencing datasets generated from our lab and other publicly available datasets, normalization and differential expression analysis (when applicable) was conducted using the edgeR package within R^143,144^. Gene detection cutoff for expression filtering before library normalization was set to a CPM > 2 (CPM = counts per million) in at least a fourth of the samples included for analysis. For DS studies, isogenic trisomic-disomic pairs were specified in source materials. For external datasets, replicates from each individual (non-DS datasets, except ES derived datasets) or specific cell-lines/subclones (DS datasets) were merged after expression filtering but prior to normalization using sumTechReps. The log2(CPM) values shown in gene count plots were calculated by the edgeR function cpm with prior.count of 2 and log2 transformation.

For the three non-DS datasets assessed, samples from each individual cell-line (numbered) were color-coded based on three tiers of *RWDD2B* expression level, providing a visual aid for how *RWDD2B* expression relates to expression levels of neighboring genes. Cell-lines were designated as having “high” *RWDD2B* expression if their log2(CPM) was greater than the 25^th^ percentile (Q1); “low” *RWDD2B* expression if this this was below the log2(CPM) value for Q1 * 0.25; and “mid” *RWDD2B* expression as anything in-between.

### Differential expression analysis for *XIST*-based silencing across clones and “selective homolog silencing (or loss)”

For differential expression analysis of our lab’s data using edgeR, the design matrix modeled the effect of dox across replicates within each clone, using a factor for clone:dox with levels for each clone with and without dox. Contrasts were set up to compute average dox response between groups of clones, assessing all *XIST*-transgenic clones (c1, c5, c5a, c7), clones silencing H1^RWD^ (c1, c5, c5a), clones silencing H2 (c7), and the difference of silencing H1^RWD^ versus H2 (i.e. selective homolog silencing; c7 vs. other clones). Because these contrasts test for differences in average responses across groups of clones while residual variance is based on between-replicate rather than between-clone variance, differences could be called significance even if they reflect changes that occur in only one or two clones, limiting confidence, especially for H2 silencing (based on only c7) and H1^RWD^ vs H2 (three vs one; imbalanced comparison). Likely due to this limitation, only highly divergent genes (*RWDD2B* and *AP001469.3*) reached significance. To confirm *RWDD2B* findings were not clonal artifacts (an oddity in c7), we analyzed prior microarray data from additional H2-silencing clones (c2, c3). Future work could improve detection by including multiple clones silencing each chr21 homolog.

The framework used above for contrasting average effects of doxycycline (dox) in different groups of *XIST*-transgenic clones can be used for controlling for the effect of dox alone in the *XIST*-transgenic groups, by contrasting with the average effect of dox in our isogenic *TET3G* expressing (TET) trisomic subclones that lack the *XIST* transgene. This approach was used to adjust for the effect of dox in *XIST*-transgenic results across all subclones, and for H1^RWD^ and H2 silencing: genes similarly affected by dox alone show reduced significance and fold change, while genes oppositely affected by dox alone show increased significance and fold change. As in the limitations mentioned above, these models do not account for variation in the dox effect for different clones; as such, dox correction was not applied when comparing the silencing of different homologs, as effects would cancel out. All analyses used voomLmFit^145^. To identify candidate *trans* effects of selective homolog silencing, an additional expression cutoff of log2(CPM) > 2 across all samples was applied after differential expression to non-chr21 genes, to remove noisy genes due to low-to-modest expression.

For our analysis of “selective homolog loss” in data from Gonzales et. al.^73^, differential expression analysis was conducted similarly to our *XIST*-transgenic data. However, as each clone lacked replication the design matrix did not include a factor for each individual clone but rather one factor for chr21 haplotype (as detailed in source supplement material) with levels for: 1) the three trisomic subclones, 2) the two subclones that lost a specific chr21 homolog (Hx), and 3) the one subclone that lost a different chr21 homolog (Hy). Contrasts were set up to compare trisomic subclones to Hx subclones, trisomic subclones to Hy subclones, or the difference between these two comparisons (i.e. the difference from losing different homologs). Due to lack of replication for each clone, and imbalance in number of clones losing different homologs, analysis of this dataset was only used to introduce the potential expansion of the approach.

### Chromosome 21 CpG methylation analysis

While described in more detail in the source publication^38^, our parental (par) trisomic DS line and two *XIST*-transgenic subclones, targeting chr21 homologs H1^RWD^ and H2 (c1 and c2 respectively), were grown with and without dox for 22 days, in duplicate cultures. Genomic DNA was extracted using PureLink Genomic DNA mini kit (Invitrogen) and bisulphite modified using the EZ DNA methylation kit (Zymo Research). Bisulphite DNA was amplified, fragmented, and hybridized to Illumina Infinium HumanMethylation450 array. Supplementary files from Illumina were used to update the locations and gene names to the current genome build (hg38).

To evaluate differences in methylation across chr21 after silencing H1^RWD^ (ΔH1) or H2 (ΔH2), average methylation values were calculated for each clone (±dox) at each individual CpG. The difference between homologs was computed as ΔH1–ΔH2. Average ΔH1–ΔH2 values were computed for CpGs grouped by gene association (nearest TSS) and further between CpG shores (when applicable), or intergenic regions between gene-associated CpGs. Two adjustments were made to these scores: 1) to down-weight scores in regions with large baseline methylation differences between non-induced clones (no dox), baseline differences were calculated for each CpG, averaged by group, rescaled between 0–1 (*scales* package), and multiplied by the average ΔH1–ΔH2 value; 2) to reduce bias from groups with few CpGs, CpG counts per group were rescaled between 1–3 and also used to weight the average ΔH1–ΔH2 values. After these adjustments, intergenic regions were excluded from analysis. The original data and additions made described above can be found in Supplement Table S5. The last 16 columns contain new additions, all other columns come from the source publication.

### Whole genome sequencing and *RWDD2B* region phasing

DNA from our parental DS iPSC line was extracted using QuickExtract (BioSearch Technologies) and sent to Azenta Life Sciences for standard Illumina whole-genome sequencing (WGS). We extracted the *RWDD2B* region (50kb upstream and downstream from *RWDD2B*) to locally phase the three chr21 homologs. As this region was small and no current R package exists to processes phasing in the presence of 3 homologs, we used the LDhap function from the R package LDlinkR^146^ to semi-manually phase heterozygous SNPs in the *RWDD2B* region. Although heterozygous SNPs were found in *RWDD2B*’s 3’ UTR in the WGS data, low RNA-seq coverage prevented their use for establishing which allele was silenced or lost in each clone. Instead, coding SNPs in the neighboring genes *LTN1* (rs2254796) and *USP16* (rs2274802), differentially present in RNA-seq data of disomic lines lacking H1^RWD^, were used to anchor phasing. To fully phase these exonic SNPs, a third coding SNP in the neighboring gene *N6TIAM1* (rs1997606) was used; however, after this, the three chr21 homologs (H1/H2/H3) could be determined by rs2254796 (G/G/A) and rs2274802 (A/T/A) for H1, H2, and H3 respectively. While rs1997606 was not used as an anchor in primary phasing rounds, it was used in the two supplemental rounds (described below) as space was available. SNPs were primarily phased in five rounds (due to input size limits for LDhap) using the CEU population from hg38 in batches of 30 (28 excluding the two exonic SNPs anchors), where 113/122 SNPs were phased. Batches were determined by descending reference read ratio without replacement.

After this, two unphased SNPs were resolved by hg38 location with matching bases and another three by switching to hg19 respectively. rs768855631 was removed, as the two different alleles were not present in the CEU population and are only found with different alleles in two individuals across all populations. SNPs used for each phasing round are listed in Supplement Table 1. As denoted in Supplement Table 1, a small handful of SNPs leveraged read coverage information for more confidently phasing. Overall, 122 heterozygous SNPs around *RWDD2B* were examined: 88 phased and in GTEx (2 non-significant), 15 phased but not in GTEx, and 19 unphased. While phasing could be improved with disomic WGS, using just trisomic cells avoids the laborious effort of generating clones that lose specific chr21 homologs.

### GTEx Data

Data files regarding *RWDD2B* eQTLs across multiple tissues were directly downloaded from GTEx (version 10). We then filtered eQTLs to contain only the heterozygous phased SNPs within our cell line. Two non-significant SNPs that were still assessed in GTEx, identified using the GTEx frozen lookup table, were also assessed.

### Graph and Figure generation

All graphs were generated in R using the ggplot2 package^147^. Diagrams were generated using BioRender.

### Data and Code availability

All code used for this experiment is available upon request. All expression data used in this study are publicly available from GEO or directly from source publications. The whole genome sequencing data from our tiromic line is pending submission to dbGap (in process).

### Material availability

The stellaris *RWDD2B* probe was made by Biosearch Technologies using the custom design tool. The sequences targeted range was exon 1 through intron 1. The specific sequences targeted by this probe batch are available upon request.

## Acknowledgements

Sources of funding are as follows: R01HD091357, R01HD094788, and R35GM122597 to J.B.L, F31HD106741A to E.C.L., F31HD095588 to J.M.

## Author Contributions

J.E.M conducted the RNA sequencing experiments described herein. E.C.L. designed and conducted all other experiments and analysis of various datasets. O.D.K provided support for RNA sequencing analysis. J.B.L., O.D.K., and E.C.L wrote the paper.

## Lead Contact

Jeanne Lawrence;

## Supplement Table Titles

**Table S1. Results from tests of differential gene expression in trisomic vs disomic organoids.**

(Excel spreadsheet Table S1)

**Table S2. Results from tests of differential gene expression based on XIST-based trisomy silencing in iPSCs.**

(Excel spreadsheet Table S2)

**Table S3. Results from tests of differential gene expression based on XIST-based trisomy silencing in iPS-derived endothelial cells.**

(Excel spreadsheet Table S3)

**Table S4. Results from tests of differential gene expression based “selective homolog silencing” in iPSCs.**

(Excel spreadsheet Table S4)

**Table S5. Chromosome 21 CpG methylation with and without XIST induction when silencing different homologs.**

(Excel spreadsheet Table S5)

Table S6. Phasing of heterozygous variants in the *RWDD2B* locus across chr21 homologs.

(Excel spreadsheet Table S6)

**Table S7. Results from tests of differential gene expression based “selective homolog silencing” in iPS-derived endothelial cells.**

(Excel spreadsheet Table S7)

Table S8. Summary of approximate *RWDD2B* expression difference between isogenic trisomic/disomic lines in Gene Expression Omnibus.

(Word doc Table S8)

---

**Supplement Figure 1:**
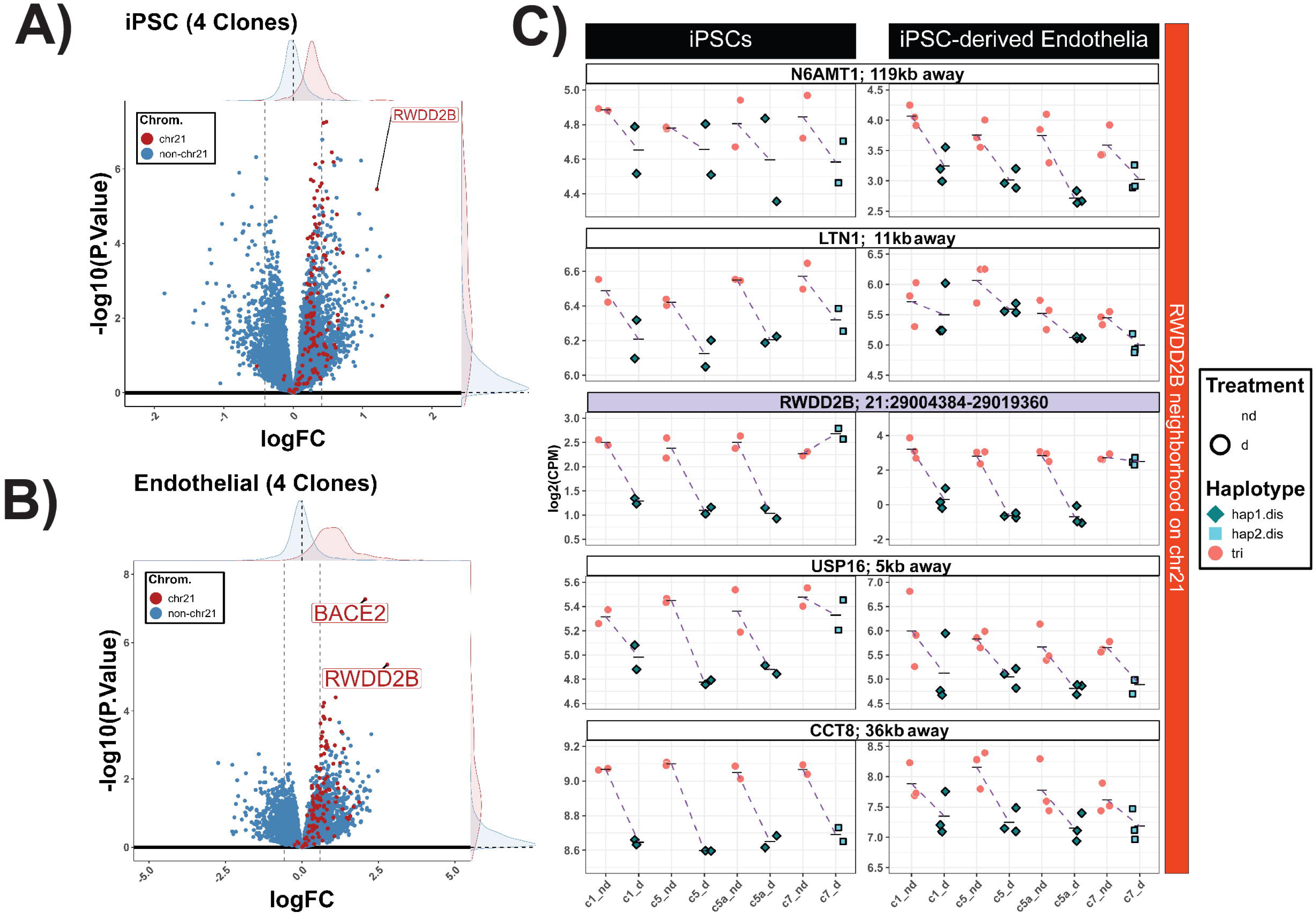
**Initial identification of RWDD2B’s outlier expression difference following trisomy silencing. A+B**) Volcano plots showing the average change due to *XIST*-based chr21 silencing across all four clones tested (regardless of which chr21 homolog was silenced) in iPSCs and iPSC-derived endothelial cells respectively. Both indicate that few other chr21 genes strongly deviate from the expected 1.5-fold increase in expression with T21 (dotted vertical line) **C)** Count plots for *RWDD2B* (purple title) and neighboring chr21 genes in both cell types indicate that expression change for *RWDD2B* depend strongly on which chr21 homolog is silenced whereas those for neighboring genes do not. Volcano plots: Red = chr21, Blue = non-chr21, Density for chr21 and non-chr21 displayed on the axis edges.

**Supplement Figure 2:**
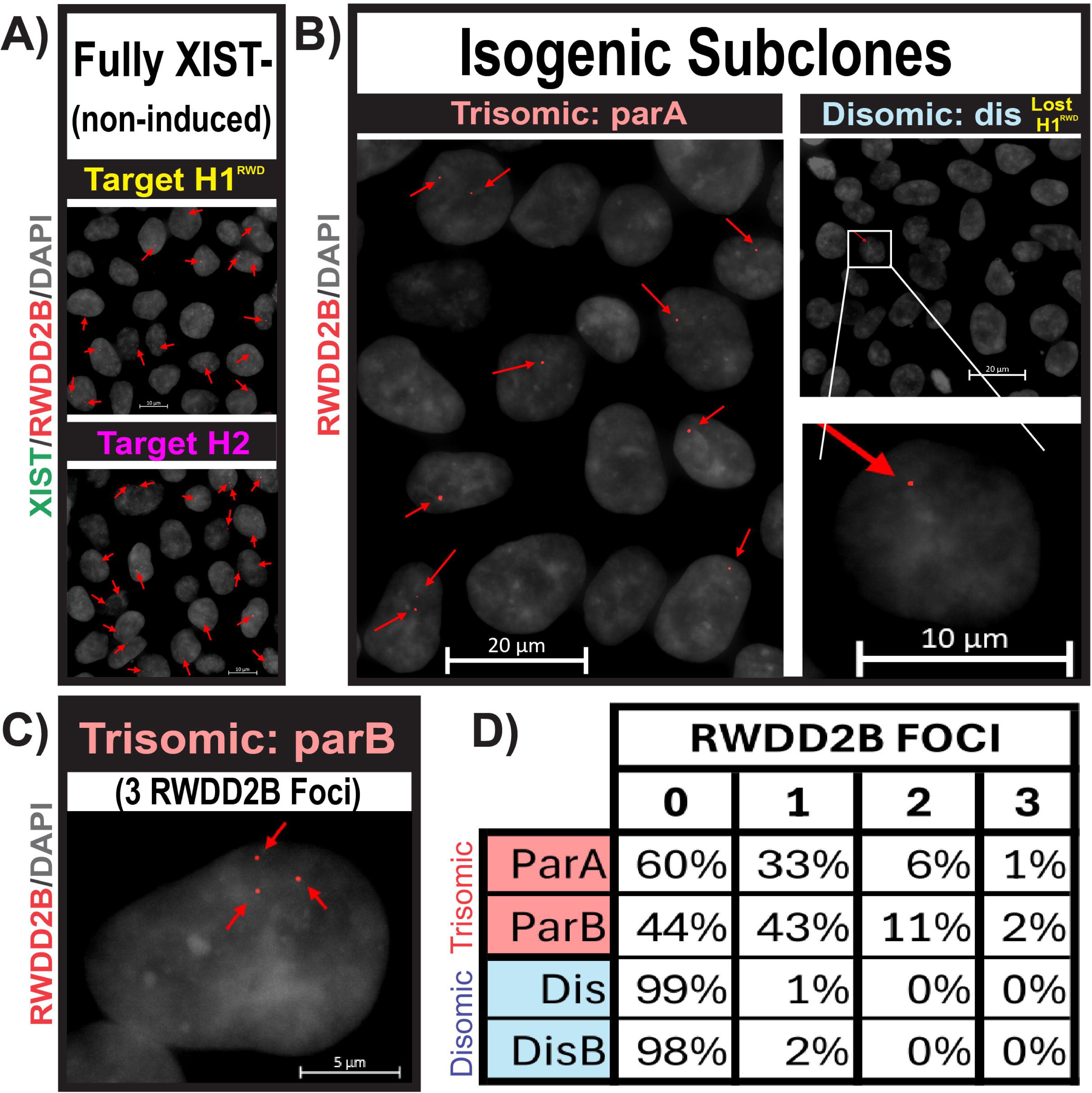
***RWDD2B* allele-specific quantification in non-induced XIST-transgenic and isogenic trisomic/disomic subclones. A)** RNA FISH for *RWDD2B* (Red) and *XIST* (Green) in non-induced XIST-transgenic cultures where in subclones that are inserted into either H1^RWD^ (c5; top) or H2 (c7; bottom) but were not induced to express *XIST* and remained trisomic. *RWDD2B* foci indicated by red arrows. **B)** <u>Left</u>) *RWDD2B* FISH in one trisomic subclone (parA) that shows primarily single *RWDD2B* foci, though some cells do have any and others have two foci. <u>Right</u>) *RWDD2B* FISH in one disomc subclone (disB), which lost H1^RWD^, indicating lack of *RWDD2B* transcription in nearly all cells; one of the few positive cells is inlayed below. **C)** While very infrequent, some trisomic cells were scored to have three clearly separated *RWDD2B* transcription foci. **D)** RNA FISH quantification in our isogenic trisomic and disomic subclones (two clones each; >200 cells quantified for each subclone), showing *RWDD2B* foci loss only with loss of H1^RWD^ and *RWDD2B* transcription foci detection rates per cell in each subclone.

**Supplement Figure 3:**
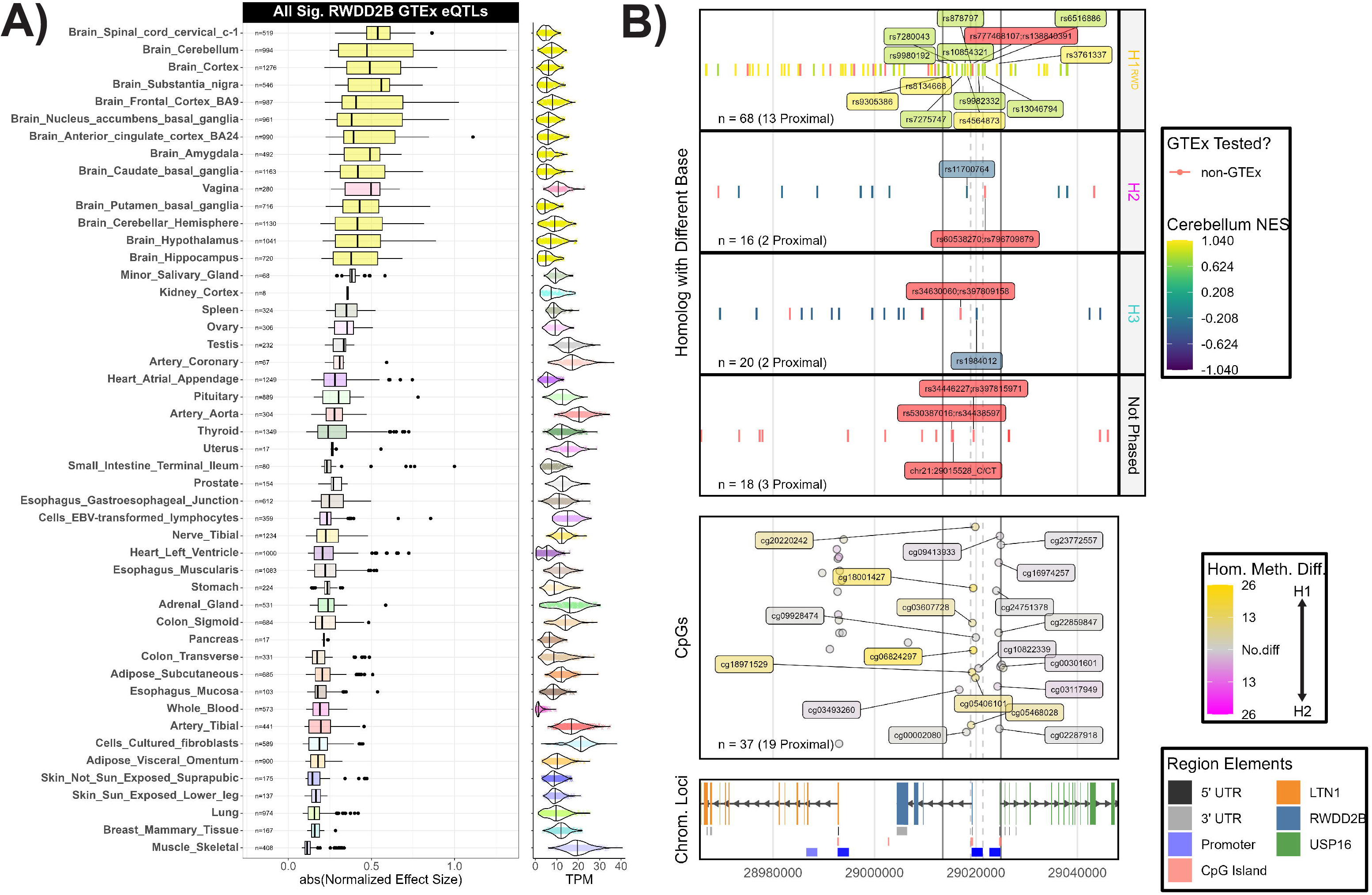
Narrowing of implicated *RWDD2B* variants from GTEx data using corresponding information from a single cell-line. **A)** GTEx data displaying the range of impacts for all significant *RWDD2B* eQTLs across tissues suggests a more modest impact and cell-type specific effect than narrowed and linked to specific chr21 homologs in our data. Right) *RWDD2B* TPM in each cell type. **B)** Narrowing of candidate variants in our cell-line in proximity to *RWDD2B*. <u>Bottom</u>) Genetic coordinates with *LTN1* (orange), *RWDD2B* (blue), and *USP16* (green) gene annotation and other regulatory elements within 50kb of *RWDD2B*. The *RWDD2B* regulatory region (Promoter + CpG Island) is indicated by vertical dotted lines and the corresponding narrowed candidate region 5.5kb up/down from this region is indicated by vertical dotted lines. <u>Middle</u>) Adjusted methylation difference between silencing of either H1^RWD^ (c1) or H2 (c2); showing when methylation increase more when silencing H1^RWD^ (yellow) vs H2 (magenta). Only CpGs near the center of the regulatory region display a strong difference. <u>Top</u>) Individual variants in our cell-line either phased to determine which chr21 homolog contained the heterozygous base (H1^RWD^, H2, or H3) or those that could not be phased (bottom). For those also found in GTEx, the effect size in the cerebellum was adjusted to reflect the ref/alt status of the heterozygous base; to display effect sizes corresponding to increased expression (yellow) vs decreased expression (purple). Phased variants not found in GTEx; with no corresponding cerebellum effect size quantified are colored red. Data indicates H1^RWD^ corresponded with higher expression while H2 and H3 correspond with lower expression. Names for specific variants and CpGs within the 5.5kb region up/down from the RWDD2B regulatory region are labeled.

**Supplement Figure 4:**
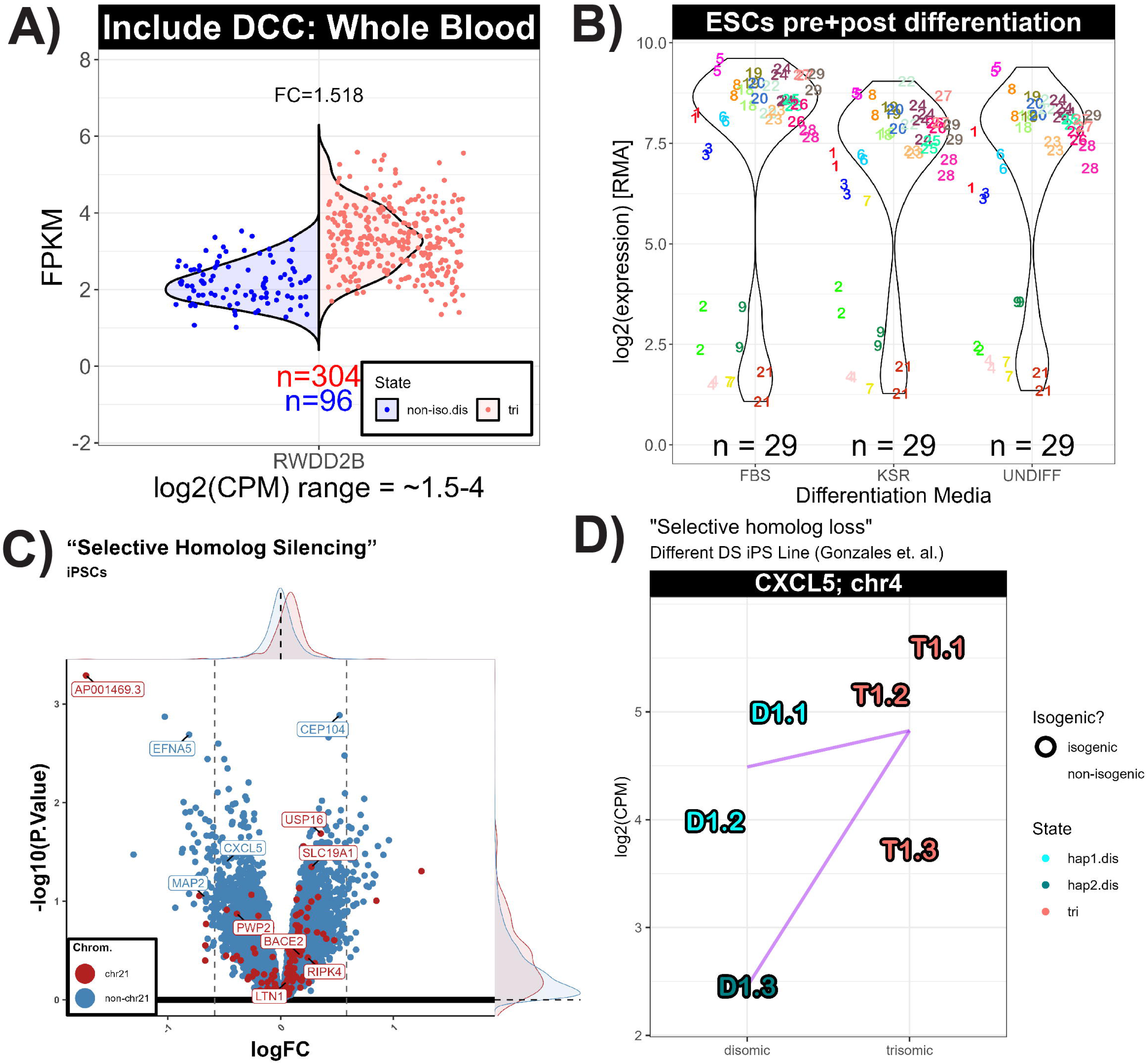
***RWDD2B* expression across many individuals in non-iPS derived samples. A)** *RWDD2B* expression distribution (in FPKM) from whole blood from ∼300 individuals with DS (tri, red) and ∼100 euploid individuals (non-iso.dis, blue) from the Include database. As suggested by our data, the whole blood samples do not suggest abnormally strong *RWDD2B* eQTLs in either population nor does any individuals (regardless of DS) show drastically less *RWDD2B* expression (as seen in other data from specific cell types). **B)** Microarray data from a study of ES cell-lines from many individuals without differentiation (UNDIFF) and following differentiation using two different media types (FBS and KSR). Each cell-line from a distinct individual is uniquely numbered and colored, with two replicates per individual. In line with the relative RMA from our iPS data (Fig. 1E; Right), specific individuals also display much lower *RWDD2B* expression in ESCs pre/post differentiation. **C)** Volcano plot comparing silencing H1^RWD^ vs H2 to identify candidate chr21 eQTLs with more modest effects than *RWDD2B* (*RWDD2B* and non-chr21 genes with log2(CPM) < 2 removed) in iPSCs. **D)** *CXCL5* expression data from Gonzales et al. showing that the subclone that lost an RWDD2B-associated chr21 homolog (dark blue) had a stronger decrease in *CXCL5* relative to isogenic trisomic subclones (red) than the two subclones that lost a different chr21 homolog (light blue). All subclones are isogenic (outlined). Volcano plots: Red = chr21, Blue = non-chr21, Density for chr21 and non-chr21 displayed on the axis edges. Count plots (D): Pink = Trisomic subclones, Disomic subclones with loss of different chr21 homolog (light vs dark blue), Purple lines connect average expression between haplotypes.

