## Supplemental Table 3 for "Selective chr21 homolog silencing reveals polymorphisms influence the epigenetic silencing and functional dosage of RWDD2B"

| **RWDD2B expression change with chr21 loss** | **# of RWDD2B expressing alleles w/ T21** | **# of RWDD2B expressing alleles w/ D21** | **PMID** | **GEO Accession** | **Cell Type(s)** |
| --- | --- | --- | --- | --- | --- |
| **Expected decrease, still expressed** (loss of high expressing allele with 1–2 high expressors remaining) | >2 | >1 | 23716668 | GSE48611 | iPSC + neurons |
|  |  |  | 38838150 | GSE208440 | cerebral organoid |
|  |  |  | 38357436 | GSE247990 | embryoid bodies + neuroectoderm |
|  |  |  | 38433139 | GSE213409 | iPSC |
|  |  |  | 29789608 | GSE110064 | B-cells |
| **No significant decrease, still expressed** (loss of non-expressing allele with 1–2 high expressors remaining) | 1 or 2 | 1 or 2 | 38245602 | GSE238115 | hematopoietic progenitors |
|  |  |  | 36313423 | GSE95552 | neural progenitors |
|  |  |  | 37792788 | GSE203257 | endothelial |
|  |  |  | **35947952** | **GSE166849** | **iPSC + endothelial** |
| **Very large decrease, essentially "off"** (loss of the only high expressing allele) | 1 | 0 | 31130512 | GSE124513 | cerebral organoid |
|  |  |  | 38848354 | N/A | iPSC + neurons |
|  |  |  | 24375627 | GSE52249 | iPSC* (mono-zygotic twins) |
|  |  |  | 29584757 | GSE101942 | iPSC |
|  |  |  | **35947952 + 36817096** | **GSE166849 + GSE222365** | **iPSC + endothelial + cerebral organoids** |

| **Columns** | **Meaning** |
| --- | --- |
| RWDD2B expression change with chr21 loss | Roughly estimated RWDD2B expression change category with loss of a single chromosome 21 in trisomy 21 cells. |
| # of RWDD2B expressing alleles w/ T21 | Estimated number of chromosome 21 homologs that highly express RWDD2B when trisomic |
| # of RWDD2B expressing alleles w/ D21 | Estimated number of chromosome 21 homologs that highly express RWDD2B when trisomic cells that became disomic by losing a single homolog |
| PMID | PubMed Identifier for each respective study |
| GEO Accession | Gene Expresssion Omnibus (GEO) Accession for each study (if available) |
| Cell Type(s) | Cell Types (or multiple cell types) examined by each study |

Additional Notes:

**Bold** studies are those conducted and published previously by the Lawrence lab.
